# Reconstruction of the human nigrostriatal pathway in vitro reveals target-dependent dopamine neuron maturation

**DOI:** 10.64898/2025.12.23.696164

**Authors:** Edoardo Sozzi, Sara Corsi, Andreas Bruzelius, Giorgio Scordo, Samuel Tavares da Silva Maraschin, Surangrat Thongkorn, Arto Heiskanen, German Ramos-Passarello, Janko Kajtez, Jenny Emnéus, Malin Parmar

## Abstract

The human nigrostriatal pathway, comprising dopaminergic neurons in the ventral midbrain (vMB) projecting to the dorsolateral striatum, is essential for motor control and selectively vulnerable in Parkinson’s disease (PD). How this circuit assembles during development and how it degenerates under pathological conditions remains poorly understood in a human context and *in vitro* models capturing its long-range connectivity and spatial organization have been lacking. Here, we introduce the *connectoid*, a compartmentalized, human stem cell-based model of the nigrostriatal pathway that integrates vMB and striatal organoids within a custom-engineered microfluidic device, confining cell bodies while guiding axonal growth, mimicking the *in vivo* topography. Functional connectivity was confirmed by retrograde rabies tracing, and optogenetic and pharmacological stimulation, while 6-hydroxydopamine-induced selective degeneration of dopamine neurons, recapitulating a key feature of PD. Additionally, single-cell transcriptomics revealed that interaction with striatal targets enhances dopaminergic neuron maturation and activates transcriptional programs linked to synaptic signaling. Thus, connectoids uniquely allow spatial segregation of regionalized organoids while preserving long-range communication, providing a scalable and physiologically relevant platform for studying human circuit assembly, selective vulnerability, and therapeutic interventions in PD.

## Introduction

Three-dimensional (3D) human brain organoids have recently emerged as a promising platform to study brain development, neuronal function, and dysfunction *in vitro* (Faravelli *et al*, 2025; Pasca *et al*, 2025). While regionalized organoids are valuable for developmental studies, disease modelling, and drug screening (Uzquiano & Arlotta, 2022), they are mainly limited by the representation of only a single brain region. To overcome this, fused organoids recapitulating two or more brain areas -referred to as assembloids -have been developed to model inter-regional processes such as cell migration and circuit formation, in both physiological and pathological conditions (Levy & Pasca, 2023; Miura *et al*, 2022). Notably, cortical assembloids generated by fusing dorsal and ventral forebrain organoids have enabled the study of aberrant interneuron migration in Timothy syndrome (Birey *et al*, 2017; Birey *et al*, 2022). Similarly, spatially arranged ventral midbrain (vMB)-striatal-cortical assembloids have been used to model dopamine (DA) neuron interactions and aspects of Parkinson’s disease (PD) with both striatal (STR) and cortical targets (Reumann *et al*, 2023; Tran *et al*, 2025). Despite these advances, assembloids do not fully recapitulate long-range neuronal connectivity, as the precise and long-range topographical organization of human axonal projections is difficult to achieve through organoid fusion alone. As an alternative, more complex systems have been developed where organoids are physically separated but connected via their projections (Kirihara *et al*, 2019; Martins-Costa *et al*, 2024; Osaki *et al*, 2024). These systems present different engineering solutions for physical compartmentalization and for providing support for axonal growth between compartments and have primarily been designed and used to study cortical developmental defects involving projection dynamics (Cullen *et al*, 2019; Kirihara *et al*., 2019; Martins-Costa *et al*., 2024; Osaki *et al*., 2024).

In this study, we introduce *the connectoid*, a human stem cell-based *in vitro* model of the nigrostriatal pathway, that enables reconstruction of long-range DA projections and their interactions with STR targets. By integrating vMB- and STR-patterned organoids into a custom polydimethylsiloxane (PDMS) device, we recapitulated the topographical organization of this pathway, which is central to motor control and selectively vulnerable in PD. Our results demonstrate that this system supports the growth of DA axon bundles, the establishment of axonal connectivity, and could be used to mirror the specific cell loss of PD. Together, these findings establish connectoids as a versatile and scalable platform for studying human-specific mechanisms of circuit formation, maturation, and pathological dysfunction. We subsequently showed that DA neurons exhibited higher expression of genes associated with neuronal maturation and showed increased spontaneous DA release when maturing in contact with their STR targets in connectoids, compared to DA neurons within vMB-vMB controls.

A defining feature of connectoids, as opposed to assembloids, is their ability to model axonal communication between spatially separated brain regions. This spatial segregation allows investigation of long-range neuronal interactions without the confounding influence of direct cell-cell proximity. To examine how such organization influences neuronal identity at the molecular level, we compared the transcriptional profiles of vMB DA neurons cultured in vMB-STR connectoids, where the two regions are physically separated and connected only via axonal projections, with those grown in vMB–STR assembloids, which permit axonal projections but lack spatial separation. Interestingly, DA neurons in connectoids exhibited upregulation of multiple genes associated with axon development and neuronal projection formation, suggesting that transcriptional programs supporting long-range connectivity are more strongly activated in the connectoid configuration and demonstrating the benefits of a model that mimics long-range connectivity.

## Results

### Generation of human vMB-STR connectoids

To recapitulate the nigrostriatal circuitry *in vitro* we first generated regionally patterned organoids of vMB and STR identity from human pluripotent stem cells (hPSCs) (Fig. 1A,B), following established protocols (Miura *et al*, 2020; Sozzi *et al*, 2022b). Neural induction was achieved in both protocols via dual SMAD inhibition, simultaneously blocking TGF-β and BMP signaling at an early stage. For vMB patterning, organoids were treated with a GSK3 inhibitor for caudalization (CHIR99021), SHH for ventralization, and FGF8 to mimic midbrain–hindbrain boundary cues. Gene expression analysis at day 15, 30 and 60 confirmed correct patterning (Figs. 1C and EV1A), and immunocytochemistry confirmed the presence of FOXA2 positive cells at day 30 (Fig. 1D), and the differentiation into TH/MAP2 positive neurons by day 60 (Fig. 1E). In STR organoids, a rostral patterning was achieved through Wnt inhibition (IWP2), with Activin A and the retinoid X receptor agonist SR11237 further refining lateral ganglionic eminence (LGE) identity. Successful patterning was confirmed through gene expression analysis at day 15, 30 and 60 (Figs. 1F and EV1A). Expression of FOXP2 and GAD67 at day 30 (Fig. 1G) and the medium spiny neurons (MSN)-specific DARPP-32 protein at day 60 (Fig. 1H) confirmed their striatal identity.

**Figure 1.**
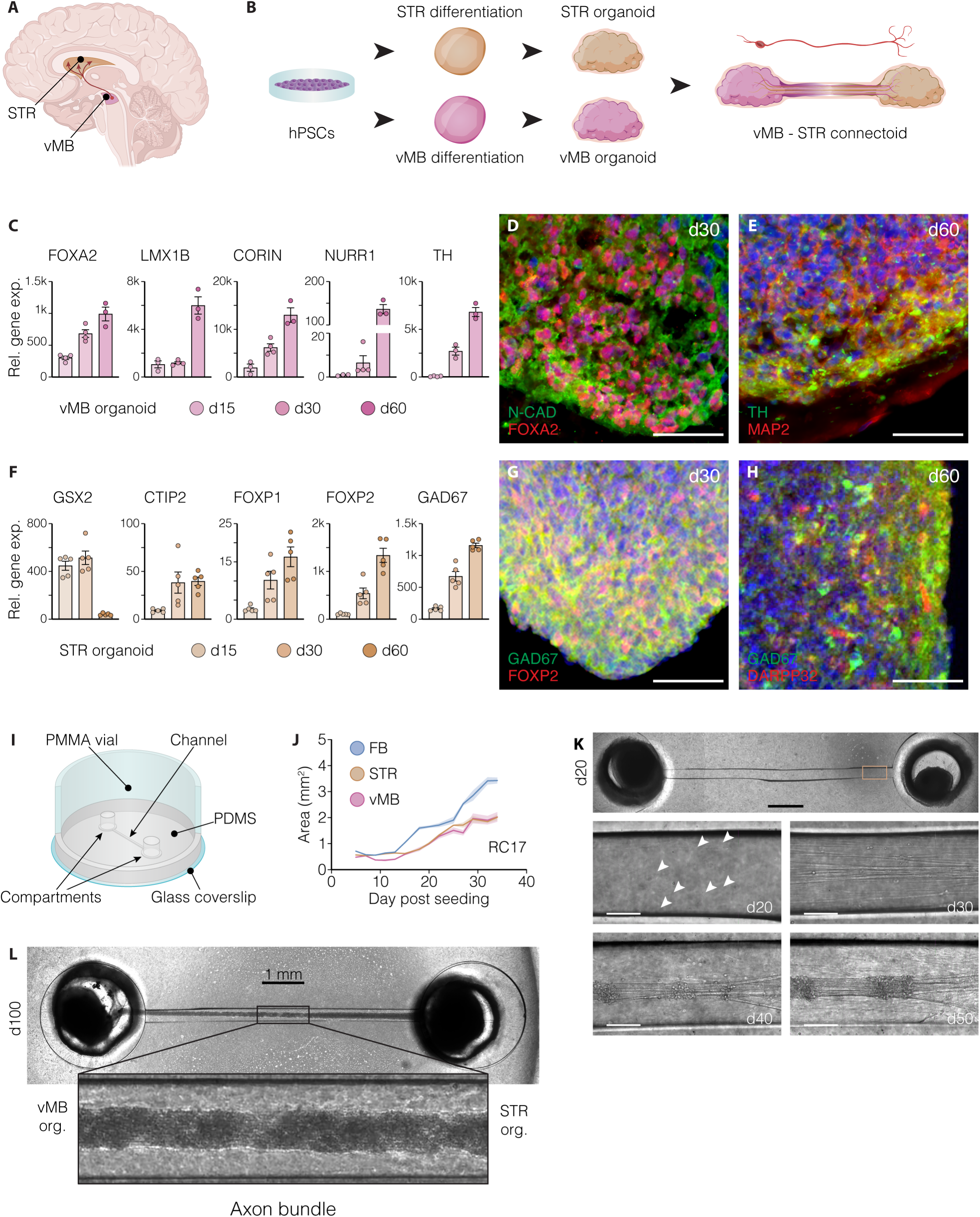
Reconstructing long-range DAergic circuitry in a microfluidic human brain model. A) Illustration of the human nigrostriatal pathway, highlighting the vMB and STR regions. B) Schematic overview of the strategy used to generate vMB-STR connectoids from hPSCs. C) Expression levels of selected markers of midbrain floorplate and DA neuron development in vMB organoids at day 15, 30, 60 of differentiation (n=4, d15; n=4, d30; n=3, d60). D,E) Immunofluorescence staining for N-CAD and FOXA2 (D, d30) and TH and MAP2 (E, d60) in vMB organoids. Scale bars, 50 μm. F) Relative gene expression over hPSCs of lateral ganglionic eminence (LGE) and striatal markers in STR organoids at day 15, 30 and 60 of differentiation (n=5). G,H) Immunohistochemistry for GAD67 and FOXP2 (G) at day 30 and GAD67 and DARPP32 (H) at day 60 of differentiation in single STR organoids. Scale bars, 50 μm. I) Schematic illustration of the custom device used for connectoid generation. J) Measurements of cross-sectional area of vMB, STR and unguided forebrain (FB) organoids across time (day 5-34 post-seeding; n=5). K) Bright field pictures of neurite outgrowth in the channel of vMB-STR connectoids at day 20, 30, 40 and 50. Scale bars, 1mm (top), 100 μm (insets). L) Representative bright field picture of a mature vMB-STR connectoid after 100 days of differentiation. Scale bar, 1 mm. Data are presented as mean ± SEM. Nuclei have been stained with DAPI.

To reconstruct the nigrostriatal pathway *in vitro*, we engineered a microfluidic device that spatially confines and guides axonal projections between physically separated organoids, thereby establishing a *connectoid* (Fig. 1I). To optimally design the size of the compartments to best suit the vMB and STR organoids, we assessed the shape, growth dynamics, and dimensions of the vMB and STR organoids from two different cell lines (RC17 and H9) over time and compared them to commonly used unguided forebrain organoids that typically are larger (Figs. 1J and EV1B-F). Based on this, we determined the optimal compartment size for vMB and STR organoids to be 2 mm in diameter. The device was thus fabricated to consist of a PDMS substrate with two 2 mm diameter compartments connected by a 9 mm long, 175 µm high, and 250 µm wide channel, which allowed axonal extension and media exchange (Figs. 1I and EV1G). Before organoid placement, both compartments and the channel were coated with 2% Matrigel to provide structural support for neuronal outgrowth. The PDMS structure supported a 17.5 mm diameter polymethyl methacrylate (PMMA) ring that formed a reservoir for culture medium, while a glass coverslip -nitric acid-treated and bonded to the PDMS by plasma treatment -sealed the device at the bottom (Figs. 1I and EV1G).

vMB and STR organoids were placed into the device after 16 days of differentiation, a stage when regional identity was already established (Figs. 1C,F, and EV1A), and the morphology remained compact and spherical, but neuronal projections had not yet emerged (Fig. EV1B,C). Already four days post-placement (day 20 of differentiation), pioneering axons started to extend from the vMB compartment into the channel (Fig. 1K). These projections progressively increased in number and complexity, forming a defined fascicle by day 40 (24 days post-placement; Fig. 1K) in a consistent and reproducible manner across multiple connectoids (Fig. EV1H). The axonal bundle continued to thicken over time, resulting in a mature and interconnected system by 3 months (Fig. 1L).

### Time-lapse imaging reveals axonal growth and degeneration dynamics of DA neurons in vMB-STR connectoids

To visualize the generation of DAergic projections in nigrostriatal connectoids over time, we employed an engineered hPSC line with a CRE recombinase knock-in into the *TH* locus that, when combined with the lentiviral delivery of a flexed GFP construct, enabled the selective labeling of TH-expressing DA neurons and their axons (Fig. 2A). Using this system, we monitored DAergic axonal growth dynamics within the connectoid channel and compartments by monitoring GFP expression using confocal imaging at 12-hour intervals, over a 10-day period (day 22–32 of differentiation) (Figs. 2B,C and EV2A,B). Quantification of TH^+^ axonal area in the vMB compartment and channel captured the progressive extension of DAergic fibers in the connectoid and its dynamics (Figs. 2B,C and EV2A,B). By day 30 (14 days post-placement), the first DAergic projections had traversed the entire 9 mm channel and reached the target STR organoids in the opposite compartment (Fig. 2B).

**Figure 2.**
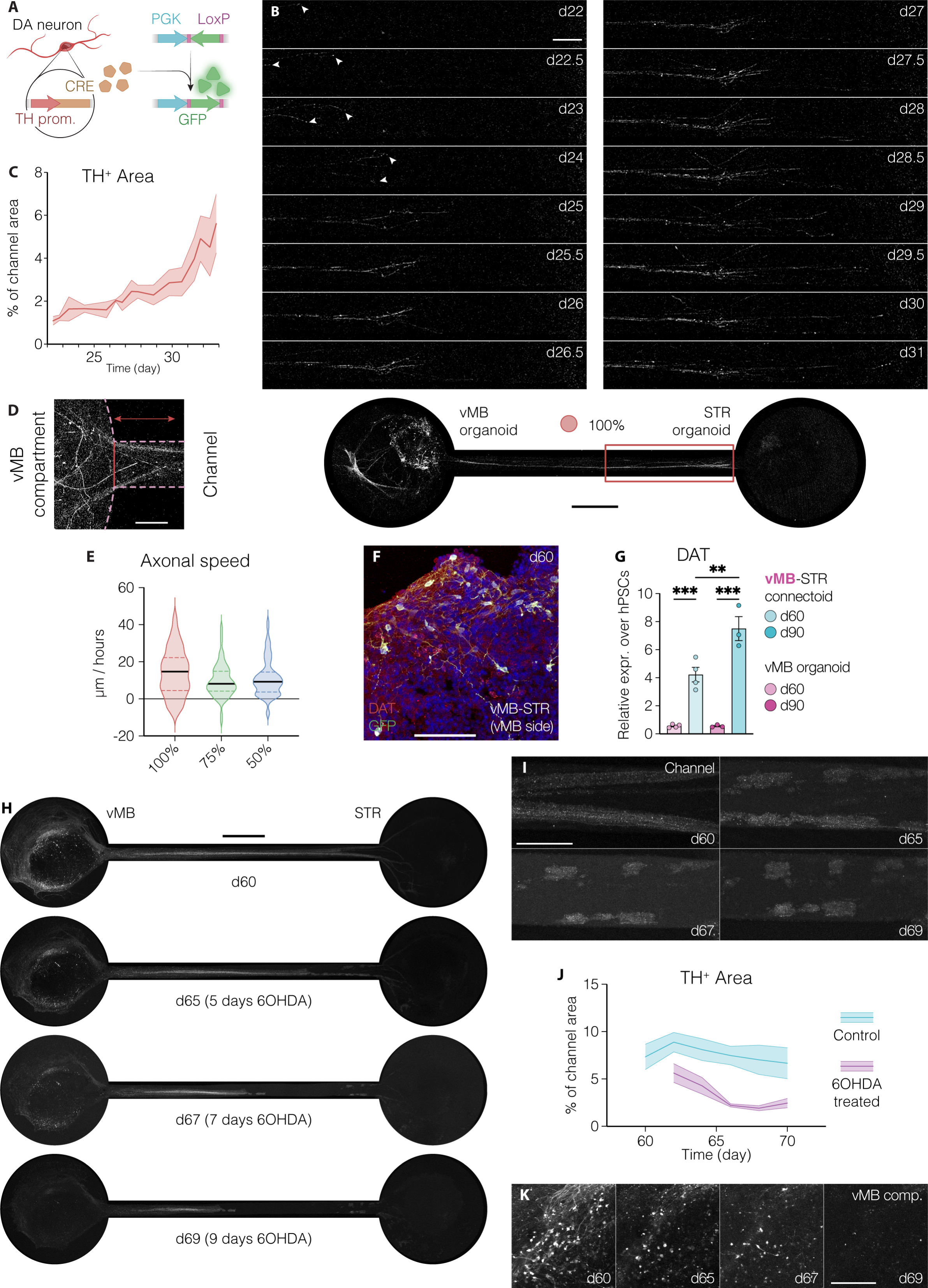
Tracking DA axonal growth and degeneration dynamics in human connectoids. A) Schematic illustration of the GFP labeling strategy to visualize TH-expressing neurons within developing vMB organoids using a TH-Cre hPSC line. B) Time-lapse images showing the outgrowth of TH/GFP⁺ neurons from day 22 to day 32 of differentiation within the channel of vMB–STR connectoids. White arrowheads indicate the growth cones of pioneer DA axons at days 22–24. Scale bars, 250 μm (top) and 1 mm (bottom). C) Quantification of TH⁺ area within the connectoid channel between days 22 and 32 of differentiation (n = 3, channel length 9 mm). D) High-magnification confocal image of the interface between the vMB compartment and the channel, used as the starting reference for neurite outgrowth measurements. E) Violin plot showing the outgrowth speed of individual DA neurites in vMB–STR connectoids with different channel lengths (100% = 9 mm, 74 measurements; 75% = 6.7 mm, 65 measurements; 50% = 4.5 mm, 37 measurements; Kruskal–Wallis test, p = 0.25). F) Immunofluorescence staining of GFP and DAT in the vMB compartment of vMB–STR connectoids at day 60. Nuclei have been stained with DAPI. G) Relative DAT expression in the vMB compartment of connectoids and single vMB organoids at days 60 and 90 (n = 4, day 60; n = 3, day 90). **p < 0.001; ***p < 0.0001; one-way ANOVA with Tukeýs test. H–I) Overview (H) and high-magnification (I) time-lapse confocal images of vMB–STR connectoids treated with 6-OHDA between days 60 and 69. Scale bars, 1 mm (H) and 250 μm (I). J) Quantification of TH⁺ area in the connectoid channel under control conditions or after 6-OHDA treatment between days 60 and 70 (n = 4, Control; n = 3, 6-OHDA; Friedman test, p=0.0005). K) Time-lapse images showing GFP⁺ DA neurons in the vMB compartment following exposure to 6-OHDA. Scale bar, 200 μm.

To further characterize our model, we investigated whether varying channel lengths influenced the dynamics of fiber growth in connectoids. For this purpose, we modified the microfluidic channel by manufacturing device variants with 75% and 50% of the original channel length, corresponding to 6.7 and 4.5 mm between compartment centers, respectively (Fig. EV2C-F). Using the channel opening on the vMB side as a reference point (Fig. 2D), we traced individual DA neuron axons based on their GFP expression and quantified the distance covered over time in the same 10-day time window considered earlier. We found that despite variability at the single-fiber level, the overall rate of axonal extension remained comparable across the devices of different sizes (mean ± SEM, 11.29 ± 0.78 µm/h) (Fig. 2E) (Kruskal–Wallis test, p = 0.25), indicating that channel length does not substantially affect projection dynamics.

A key feature of PD is the loss of DA neurons in the vMB, which is replicated in rodents with the 6-OHDA-based model of PD (Ungerstedt & Arbuthnott, 1970). The 6-OHDA toxin is internalized specifically by DA neurons via the dopamine transporter (DAT), leading to disruption of the mitochondrial chain and reactive oxygen species production, ultimately leading to neurodegeneration. We confirmed the presence of DAT and its higher expression in the connectoids compared to vMB single organoids (Fig. 2F,G), suggesting that the DA neurons in connectoids are sensitive to 6-OHDA like their *in vivo* counterparts. To test this, we exposed vMB–STR connectoids to 400 µM 6-OHDA treatment at day 60, and we monitored the presence of GFP^+^ (TH^+^) axonal projections over 9 days using the imaging setup previously described (Fig. 2H). A pronounced reduction in TH^+^ axonal area was observed already after 5 days of treatment, accompanied by ruptures in the axon bundle (Friedman test, p=0.0005) (Figs. 2H-J and EV2G). Similarly, TH^+^ axons in the vMB compartment were lost with a comparable dynamic (Friedman test, p=0.0006) (Figs. 2K and EV2H,I). Ten days after treatment, TH^+^ axons were nearly completely lost, effectively mimicking the degeneration of *substantia nigra* DA neurons in PD and the resulting disruption of the nigrostriatal pathway *in vitro* (Figs. 2H–K and EV2G–I).

Together, these data demonstrate that our system successfully recapitulates the initial long-range projections of DA neurons within the vMB compartment and their functional connections with STR target cells, effectively modeling the formation of the nigrostriatal pathway within the custom-made device. Moreover, the device can reproduce the DA degeneration characteristic of PD, providing a platform to study disease mechanisms in a human context.

### Nigrostriatal connectoids develop synaptic connections that respond to pharmacological stimuli

To directly assess the formation of synaptic contacts between vMB and STR cells in our model, we employed a modified retrograde rabies-based tracing system, Δrab (Wickersham *et al*, 2007). In Δrab, the glycoprotein (GP) needed for transsynaptic spread has been replaced with a mCherry protein, allowing the tracking of the viral spread retrogradely across one synapse and can thus be used to identify first-order synapses of vMB neurons connected to STR organoids (Fig. 3A). Before placement in the connectoid, STR organoids were transduced with a lentiviral tracing vector expressing TVA (avian receptor which allows selective entry of the rabies virus to the targeted human cells), alongside the expression of nuclear GFP (ncGFP) marking the targeted cell population, as well as the GP required for transsynaptic viral spread (Fig. 3A,B). After addition of Δrab virus on day 63, cells in the STR connectoids containing the tracing vector started to express mCherry in addition to GFP indicating successful infection (Fig. 3B). As these cells also contain GP necessary for transsynaptic spread, Δrab is retrogradely transmitted to presynaptic neurons forming synaptic contacts with the initial population and these input neurons start to express mCherry but not GFP (Fig. 3B,C). Starting from day 70, mCherry-positive fibers were detected throughout the length of the microfluidic channel, as well as in somas in the vMB compartment, indicating axonal connectivity between the regionalized organoids (Fig. 3C,D). In one example, we traced a complete projection between a vMB neuron and its synaptic partner in the STR organoid, spanning over 6 mm from soma to soma (Fig. 3E). The mCherry^+^ cells within the vMB compartment showed no ncGFP signal, confirming that its expression was due to the retrograde transport and not from the direct transduction of the TVA-containing vector. To further confirm the specificity of this system, the same vector construct lacking the GP was delivered to the STR organoids and treated with the rabies virus in parallel as a negative control. In this group, we did not detect any mCherry^+^/GFP^−^ cells in either STR or vMB compartment, confirming that viral spread depends strictly on GP-mediated transsynaptic transfer (Fig. EV3A). Together, these results confirm that vMB-derived DA neurons form synaptic connections with STR targets. To further assess DA synapse function, we implemented two complementary stimulation approaches. First, we combined the *TH*-Cre reporter line with a flexed ChR2 construct, enabling selective optogenetic activation of *TH*-expressing DA neurons with 470 nm blue light (Fig. 3F). Second, we applied pharmacological stimulation using amphetamine, which enters presynaptic terminals via DAT and promotes DA release into the synaptic cleft (Fig. 3G). If functional synapses are generated in the connectoids, both strategies should drive postsynaptic transcription of immediate-early genes such as cFOS via activation of DA receptors D1R (Fig. 3F,G). Indeed, we observed that optogenetic stimulation induced a 3-fold increase in cFOS-positive STR neurons 1.5 hours after a 10-minute light exposure (one-way ANOVA, p=0.006) (Fig. 3H,I,K). Similarly, treatment with 20 µM D-amphetamine produced a 2.4-fold increase in cFOS-positive cells relative to baseline (one-way ANOVA, p=0.04) (Fig. 3J,K). Notably, no cFOS induction was observed in connectoids with severed axonal bundles, confirming that activation required intact DA projections between the two compartments (one-way ANOVA, p=0.72) (Figs. 3J and EV3B,C).

**Figure 3.**
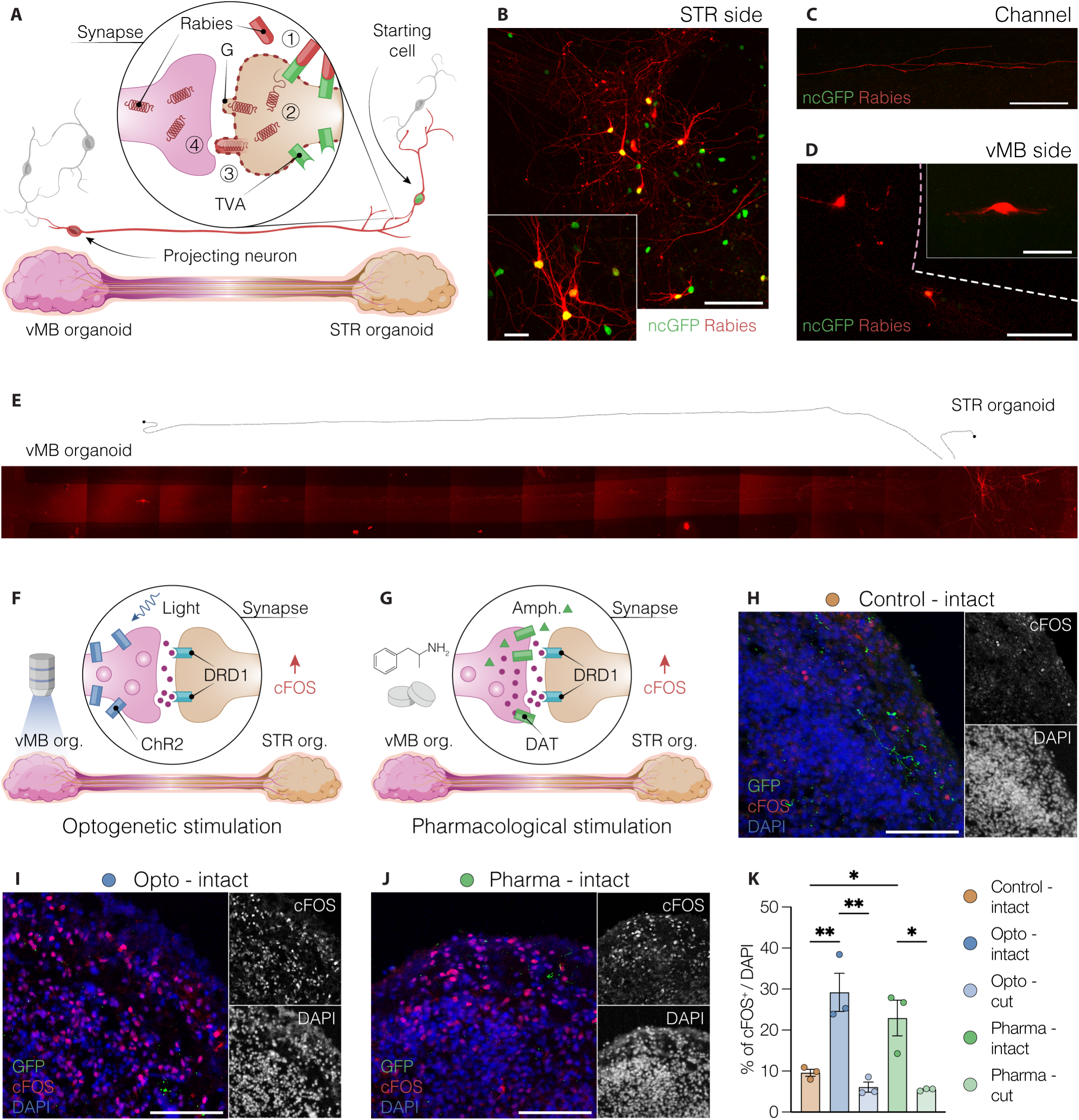
Functional DAergic axonal connectivity in nigrostriatal connectoids. A) Schematic illustration of the rabies-based tracing strategy used to identify neurons in synaptic contact. STR organoids were transduced with a TVA/GP/ncGFP construct at day 7, and first-order connecting neurons were traced using an EnvA-pseudotyped ΔG-rabies vector carrying mCherry. B–D) Confocal images of traced neurons in the STR compartment (B), channel (C), and vMB compartment (D) of nigrostriatal connectoids at day 70. Scale bars, 100 μm (B–D), 25 μm (B, inset), and 50 μm (D, inset). E) Overview and reconstruction of a single mCherry⁺ neuron projecting from the vMB organoid to the STR target in a day 70 connectoid. F, G) Schematics of the optogenetic (F) and pharmacological (G) approaches used to stimulate DA synapses in vMB–STR connectoids. H–J) Confocal images of GFP, cFOS, and DAPI immunostaining in connectoids under control conditions (H), or following stimulation with 470 nm blue light (I) or 20 μM D-amphetamine (J) at day 95. Corresponding single-channel images of cFOS and DAPI used for quantification are shown on the side. Scale bars, 100 μm. K) Quantification of cFOS⁺ nuclei relative to total DAPI⁺ nuclei in the STR compartment of nigrostriatal connectoids at day 95 across experimental conditions (n = 3). *p < 0.05, **p < 0.001; one-way ANOVA with Tukeýs test.

Collectively, these findings indicate that the DA neurons make synaptic contacts with the striatal target cells, and that DA synaptic activity in connectoids can be stimulated by both optogenetic and pharmacological approaches and the subsequent response can be recorded in the STR target cells, providing a functional framework for circuit-level investigations.

### Striatal target interactions promote maturation of midbrain DA neurons *in vitro*

Next, we assessed the DA neuron specification and maturation at the molecular level and how this is controlled by connectivity. First, we confirmed that vMB and STR organoids maintained their regional identities when cultured for an extended period of time in the connectoid devices (Figs. 4A–C and EV4A). We found that, as for single organoids, vMB-patterned cells in connectoids expressed FOXA2 at day 30, while co-expression of TH, MAP2, and NEUN emerged at later stages (day 60–90), consistent with a mature DA neuron identity (Figs. 4A–C and EV4A). In the STR compartment, staining for CTIP2, GAD67, and DARPP32 confirmed the presence of medium spiny neurons (MSNs) after 2-3 months of differentiation, following a similar timeline as for single STR organoids (Fig. EV4B–C). At this stage, the emergence of GABAergic fibers extending from the STR compartment toward the vMB side was also observed (Fig. EV4D). Gene expression profiles of vMB organoids at days 30, 60, and 90 compared to the corresponding compartments in vMB–STR and vMB–vMB control connectoids showed that floor plate and midbrain DA markers such as *TH*, *CORIN*, *EN1*, *LMX1A*, *LMX1B* and *SOX6* were consistently expressed (Fig. 4D).

**Figure 4.**
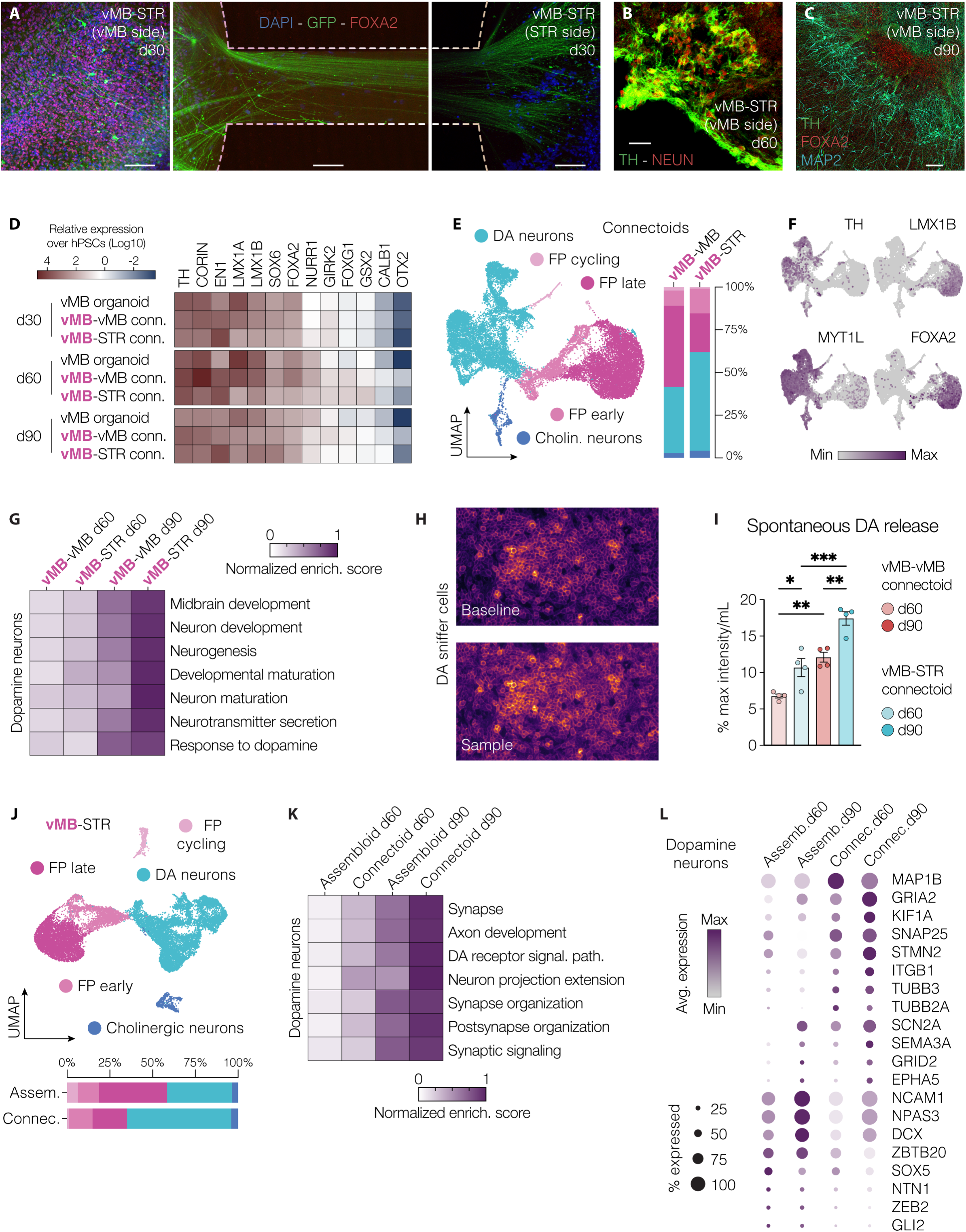
Transcriptional maturation of long-range projecting DA neurons. A) Immunofluorescence images of GFP and FOXA2 in the vMB compartment (left), channel (center), and STR compartment (right) of vMB–STR connectoids at day 30. Scale bar, 100 μm. B, C) Immunofluorescence staining of TH/NEUN (B) and TH/FOXA2/MAP2 (C) in the vMB compartment of connectoids at day 60 (B) and day 90 (C). Scale bars: B, 50 μm; C, 100 μm. D) Heatmap showing gene expression relative to hPSCs (log_10_ fold change over hPSCs) in vMB organoids at days 30, 60, and 90 under different maturation conditions (n = 3–8; outliers removed using the ROUT method, Q = 1%). E) UMAP visualization and bar plot illustrating the relative cellular composition of vMB organoids in connectoids with either STR or vMB organoids in the opposite compartment at days 60 and 90 (n = 3). FP, floor plate; Cholin, cholinergic. F) Feature plots showing the expression of key DA neuron and floor plate markers within the vMB compartment dataset. G) Heatmap displaying normalized enrichment scores for gene ontology terms associated with neuronal development and maturation within the DA neuron cluster. H) Live fluorescence imaging of GRAB_DA2M_ sniffer cells at baseline (top) and during their response to spontaneously released DA (bottom). I) Quantification of spontaneously released DA from vMB–vMB and vMB–STR connectoids at days 60 and 90 (n = 4), measured using GRAB-DA2M sniffer cells. Raw values were normalized to sampling volume and number of vMB organoids. *p < 0.05, **p < 0.001, ***p < 0.0001; one-way ANOVA with Tukey’s post hoc test. J) Relative cellular composition of vMB organoids in vMB–STR assembloids and connectoids at days 60 and 90 (n = 3). K) Gene set enrichment scores for Gene Ontology (GO) terms associated with axonogenesis and synaptic communication in DA neurons derived from assembloids and connectoids. L) Dot plot showing expression levels of selected differentially expressed genes between DA neurons derived from vMB-STR assembloids and connectoids at day 60 and 90.

To assess the effect of axonal contact with striatal targets on DA neuron maturation at the molecular level, we performed single-nuclei RNA sequencing on vMB organoids cultured as connectoids with either STR organoids or a second vMB organoid as control. At both day 60 and day 90, a diversity of cell populations was identified in the vMB compartment, including multiple classes of floor plate-derived progenitor cells (cycling: *SOX2*, *GLI3*, *CENPK*, ∼1.62%; early: *NES*, *HES1*, *VIM*, ∼11.33%; late: *NTN1*, *LMX1B*, *CORIN*, ∼36.70%), DA neurons (*TH*, *EN1*, *KCNJ6*/GIRK2, ∼47.02%), and a small fraction of cholinergic neurons (*SLC5A7*, *ISL1*, *CHAT*, ∼3.33%) (Figs. 4E,F and EV4E–G), in line with previous observations (Fiorenzano *et al*, 2021b). High-resolution analysis of midbrain DA neurons enabled a direct comparison of their transcriptomic profiles between the two maturation conditions. Notably, gene set enrichment analysis (GSEA) revealed that DA neurons in vMB–STR connectoids exhibited higher expression of genes associated with *neuronal maturation*, *midbrain development*, and *neurotransmitter secretion* compared to vMB–vMB controls (Fig. 4G). To validate these transcriptional findings, we also investigated the ability of spontaneous DA release from the vMB side within the connectoid system at both day 60 and day 90 by utilizing a microscopy-based detection system using sniffer cells, HEK cells expressing the fluorescent DA sensor GRAB-DA2M (Klein Herenbrink *et al*, 2022; Sun *et al*, 2018). The sniffer cells have been validated to show an increase in fluorescent intensity in a dose-dependent manner when exposed to DA concentrations between 100 pM and 10 nM (Kajtez *et al*, 2025; Rezaei *et al*, 2023). With this system in place, we collected media from vMB-STR connectoids at day 60 and day 90, alongside vMB-vMB connectoids as a positive control. The collected media were subsequently added onto sniffer cells to measure the DA content (Fig. 4H). This analysis revealed significantly higher levels of spontaneous DA release in vMB–STR connectoids compared to vMB-vMB controls (Fig. 4I; one-way ANOVA with Tukeýs test, day 60, p = 0.0319; day 90, p = 0.004). Together, these data suggest that interactions with target STR organoids promote both transcriptional (Fig. 4G) and functional (Fig. 4H,I) maturation of vMB-derived DA neurons in connectoids.

A distinctive feature of connectoids compared to assembloids is that they uniquely allow for studying axonal interactions while separating out the spatial proximity of two organoids. We therefore compared whether the transcriptional changes in midbrain DA neurons cultured in vMB-STR connectoids (where the cells are physically separated and only connected via axonal projections) differed from when the same midbrain DA neurons were grown in vMB–STR assembloids with axonal projections but no spatial separation. While connectoids enable independent manipulation and analysis of the two compartments, mature cells originated from vMB and STR organoids cannot be separated as easily in fused assembloids. We therefore performed this experiment using vMB and STR organoids from two distinct hPSC lines (RC17 and H9, respectively), leveraging their genetic differences to subsequently separate the data bioinformatically and restrict the comparison to vMB-derived cells across the two models. This strategy also confirmed the high purity of sample preparation in connectoids, with only 0.52% of STR-derived cells detected in the vMB compartment (Fig. EV4H), which is within the expected error margin for genetic demultiplexing methods. As in the previous dataset, the same cell types were identified, and DA neurons were subset for further comparisons across models (Figs. 4J and EV4I). Interestingly, DA neurons from connectoids and assembloids showed differential expression of multiple genes linked to *axon development* and *neuronal projections*, including *TUBA1A*, *BEX2*, *VIM*, and *IFT20*, suggesting the activation of transcriptional programs involved in long-range connectivity was enhanced in connectoids (Figs. 4K,L and EV4J). Moreover, connectoid-derived DA neurons were enriched in genes associated with synaptic compartments, signaling, and organization—such as *SNAP25*, *GRIA2*, *SCN2A*, and *GRID2*—suggesting enhanced synaptic function in long-range DA neurons (Figs. 4K,L and EV4J,K). By contrast, assembloid-derived DA neurons were enriched in *NCAM1*, a mediator of cell–cell adhesion, as well as markers of neuronal immaturity, including *NFIA*, *HES4*, *DCX*, and *NTN1* (Figs. 4L and EV4J,K).

## Discussion

Neurodegenerative disorders such as PD are increasingly viewed as disorders of neural circuit dysfunction, in which progressive disruption of long-range connectivity drives their defining motor and cognitive symptoms (McGregor & Nelson, 2019). In PD, this manifests with the selective loss of DA neurons in the vMB, particularly in the *substantia nigra pars compacta*, which project to the dorsolateral STR via the nigrostriatal pathway (Bjorklund & Dunnett, 2007; Garritsen *et al*, 2023). To date, most experimental studies of the development and degeneration of the nigrostriatal circuitry have come from animal models, where the loss of DA neurons is induced by neurotoxic agents such as 6-OHDA, the meperidine analog MPTP, or via genetic manipulation (Dawson *et al*, 2010; Fisher & Bannerman, 2019). *In vitro* models of the human nigrostriatal pathway remain limited, and existing systems do not fully capture the complexity of region-specific interactions and long-range connectivity (Di Lullo & Kriegstein, 2017; Fiorenzano *et al*, 2021c; Kelava & Lancaster, 2016).

The connectoids, as described here, successfully replicate the nigrostriatal pathway and provide a compartmentalized architecture that spatially separates the two brain regions, allowing independent manipulation and analysis of distinct biological components. Although the chosen distance between compartments does not reproduce the full length of the human nigrostriatal tract (Kordower *et al*, 2013), it offers a practical balance between the protracted timeline of human brain development and experimental accessibility, enabling systematic interrogation of synaptic functions and innervation dynamics *in vitro*. This spatial control permitted real-time monitoring of axonal dynamics, and snRNA-seq showed that the DA neurons display enhanced expression of genes associated with axonogenesis and neuronal projections.

Although connectoids, like most organoid-based systems, do not yet contain internal vascularization or microglia components that are highly relevant to both physiology and pathology (Fiorenzano *et al*, 2025), the absence of these elements can in fact be advantageous. By enabling the study of a more defined and “separated” neuronal system, connectoids offer a unique opportunity to isolate specific cellular contributions and dissect underlying mechanisms relating to circuitry with greater precision. Emerging bioengineering strategies, including the incorporation of multielectrode arrays, DA detection systems, vascularized cultures, and immune cells, are expected to further expand usability for functional and pathophysiological studies. Importantly, our findings demonstrate that even in their current configuration, nigrostriatal connectoids provide a robust and accessible platform for modeling progressive DA neuron loss following toxin exposure, thereby enabling future studies of key PD-related features in a human context. Because the devices can be fabricated using standard methods, they can be broadly implementable in laboratories without specialized infrastructure or expertise. Moreover, rabies-based tracing and cFOS induction, together with spontaneous DA release, confirmed functional axonal connectivity within the system. Finally, transcriptomic analyses revealed the upregulation of gene networks involved in neuronal maturation, neurotransmitter release, and synaptic function in the presence of STR targets, highlighting the capability of connectoids to model target-dependent maturation of DA neurons.

In summary, these findings highlight connectoids as a robust, modular platform for reconstructing and studying human long-range neural circuits *in vitro*, advancing our understanding of DA neuron maturation, circuit integration, and repair (Bjorklund & Parmar, 2021; Moriarty *et al*, 2022). Connectoids are also well suited for perturbation studies, both genetic and pharmacological, as demonstrated by the amphetamine treatment and the resulting postsynaptic activation of STR neurons. Beyond PD, the modularity of the system has broad applicability to modeling other human brain circuits, such as cortico-striatal or thalamo-cortical pathways, with only minor modifications such as adjustment of compartment size and number. Together, these features position connectoids as a powerful experimental platform for dissecting circuit-level mechanisms in health and disease.

## Materials and methods

### Human pluripotent stem cell culture

Undifferentiated hPSC lines RC17 (Roslin Cells, cat. no. hPSCreg RCe021-A), H9 (WiCell, cat. no. hPSCreg WAe009-A), and TH-Cre knock-in line (clone B2) (Fiorenzano *et al*, 2021a) were maintained on 6-well plates (Sarstedt, cat. no. 83.3920) coated with recombinant laminin-521 (Biolamina, 0.5 μg/cm^2^ in DPBS+Mg^2+^/Ca^2+^) and cultured in iPS Brew XF medium (Miltenyi Biotec, GMP-grade). At 70-90% confluence, cells were passaged using EDTA (0.5 mM for 7 min at 37 °C) and reseeded at 5k-10k cells/cm² in iPS Brew supplemented with 10 μM ROCK inhibitor (Y-27632, Miltenyi, cat. no. 130-106-538) for the first 24 hours. Unguided forebrain organoids used as controls for morphological assessments were generated according to Sozzi et al. Frontiers 2022 (Sozzi *et al*, 2022a).

### Ventral midbrain organoid differentiation

vMB-patterned organoids were generated following the protocol described in Sozzi et al. Current Protocols 2022 (Sozzi *et al*., 2022b). Briefly, cell aggregates were generated by seeding 8000 hPSCs per well in U-bottom 96-well plates (Corning, cat. no. 7007) in iPS-Brew with 10 μM Y-27632. Differentiation medium (1:1 DMEM/F12 and Neurobasal) was supplemented with N2 (1:100), SB431542 (10 μM), rhNoggin (100 ng/mL), SHH-C24II (300 ng/mL), and CHIR99021 (RC17 and TH-Cre line, 1.5 μM; H9, 0.9 μM), together with glutamine (2 mM), MEM-NEAA (ThermoFisher, 11140050), penicillin-streptomycin (20 U/ml), and 2-mercaptoethanol (50 μM). From day 8, aggregates were instead exposed to FGF-8b (100 ng/mL). At day 12, developing vMB organoids were cultured in Neurobasal with B27 (–vitamin A, 1:50), BDNF (20 ng/mL), and L-ascorbic acid (200 μM) in addition to FGF-8b. From day 16, vMB organoids were either transferred into connectoid devices with 2% Matrigel (Corning, cat. no. 354277) or embedded in 30 μL Matrigel droplets for long-term culture. Terminal differentiation medium containing Neurobasal with B27 (–vitamin A), BDNF (20 ng/mL), L-ascorbic acid (200 μM), db-cAMP (500 μM), GDNF (10 ng/mL), and DAPT (1 μM) was used for cultures up to 4 months after the beginning of differentiation.

### Striatal organoid differentiation

STR organoids were generated following the protocol described in Miura et al., Nature Biotech 2020, with minor modifications (Miura *et al*., 2020). Briefly, hPSCs were seeded in 96-well U-bottom plates as described above and cultured in differentiation medium (1:1 DMEM/F12 and Neurobasal, supplemented with 1:100 N2) containing SB431542 (10 μM, Miltenyi, cat. no. 130-106-543) and rhNoggin (200 ng/mL, Miltenyi, cat. no. 130-103-456) for the first 6 days. From day 6 to day 16, cells were maintained in Neurobasal medium supplemented with IWP-2 (2.5 μM, Selleckchem, cat. no. S7085) and Activin A (100 ng/mL, Peprotech, cat. no. 120-14E) to induce rostral fate. Starting on day 12, the retinoid X receptor agonist SR11237 (100 nM, Tocris, cat. no. 3411) was also added. Similarly to vMB organoids, STR organoids were cultured with terminal differentiation medium from day 16, ensuring compatibility between protocols in all assembloid and connectoid experiments.

### Connectoid formation

Microfluidic devices were fabricated using photolithography and replica molding. The master mold was prepared with SU8 2075 epoxy-based photoresist, exposed to ultraviolet light (500 mJ/cm² for 15 seconds), and subjected to pre- and post-exposure baking steps of 6 and 10 hours, respectively, at 50 °C. Development was performed in propylene glycol methyl ether acetate (PGMEA) for 10 minutes, followed by an additional 15-hour bake at 90 °C, yielding a master mold with a thickness of 175 μm. To facilitate Poly(dimethylsiloxane) (PDMS) release, the mold was silanized using trichloromethyl silane (1:10 dilution in toluene). PDMS was then cast onto the mold to replicate the microstructures by using a 10:1 mixture of Sylgard 184 silicone elastomer base and curing agent. Each PDMS replica (ca. 2 mm thick) was treated with oxygen plasma to enable bonding of a nitric acid–treated glass coverslip to serve as the bottom. A CNC (Computer Numerical Control) milled polymethyl methacrylate (PMMA) reservoir, designed to contain cell culture medium, was attached to the upper side of each bonded PDMS replica using freshly prepared PDMS (1:10 mixture of base and curing agent). Each device contained two organoid compartments (2 mm in diameter) connected by a 9 mm-long microchannel (250 μm × 175 μm) (Fig. 1I and EV1G). At least 24 hours prior to organoid placement, PDMS devices were sterilized with ethanol and UV light, then pre-coated with 2% Matrigel in DMEM/F12.

### RNA extraction and qRT-PCR

Total RNA extraction from vMB and STR organoids was achieved with 350 μL of RTL lysis buffer and the RNeasy Micro Kit (Qiagen). RNA quantity and integrity were assessed with a NanoDrop 2000 spectrophotometer (Thermo Fisher Scientific), ensuring a 260/280 nm ratio ≥ 2. Reverse transcription was performed with the Maxima First Strand cDNA Synthesis Kit for qRT-PCR (Thermo Fisher Scientific). Each qRT-PCR reaction contained 1 μL cDNA, 4 μL primers (0.95 μM, Integrated DNA Technologies; sequences in Table 1), and 5 μL LightCycler 480 SYBR Green Master (Roche) in 384-well plates, prepared using a Bravo Liquid Handling Platform (Agilent). Reactions were run on a LightCycler 480 II instrument (Roche) with a 40-cycle two-step protocol (95 °C, 30 s; 60 °C, 1 min). Expression was calculated with the ΔΔCT method from three technical replicates per sample, normalized to ACTB and GAPDH, and presented as fold change relative to undifferentiated hPSCs.

**Table 1.**
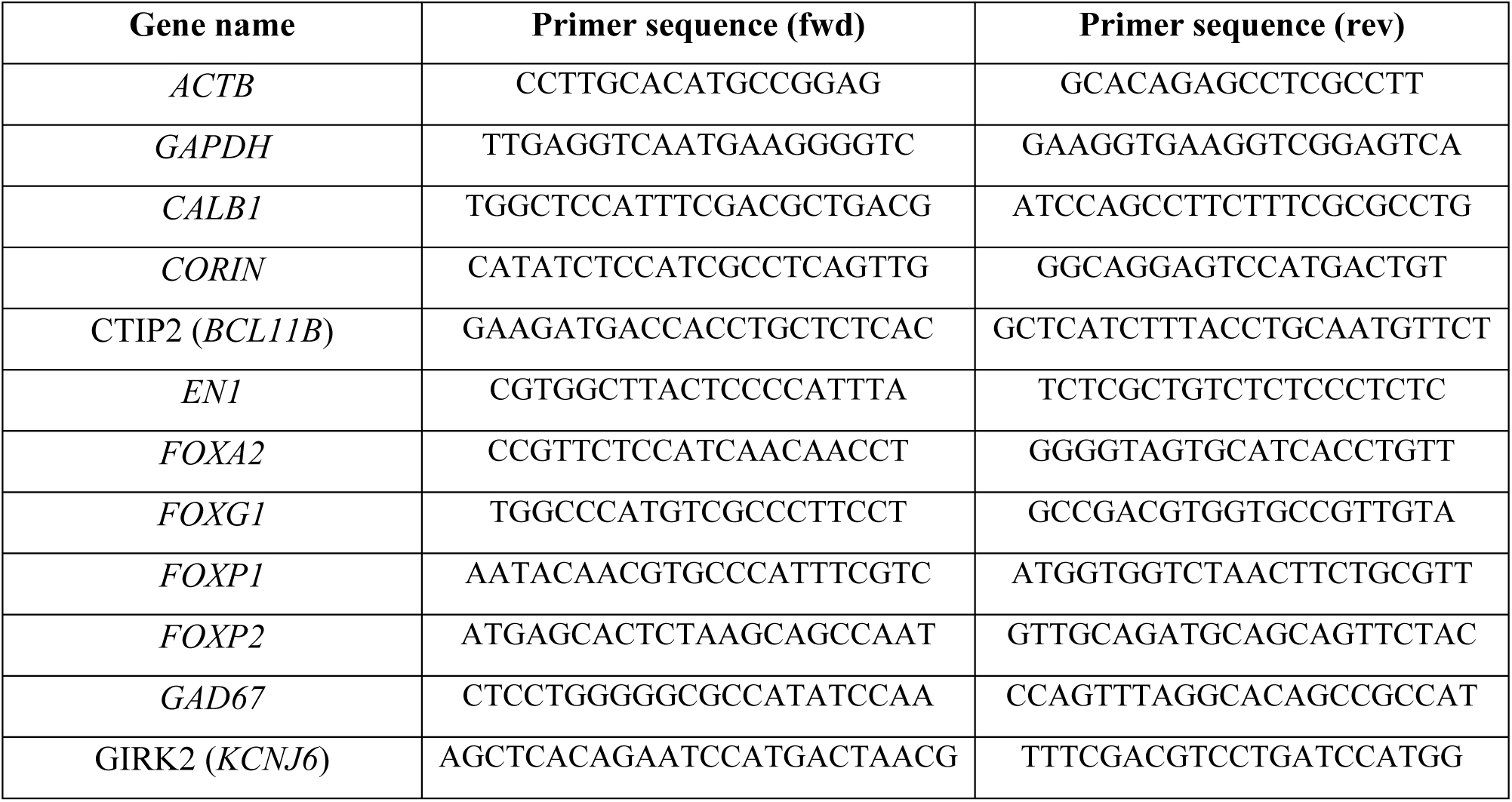
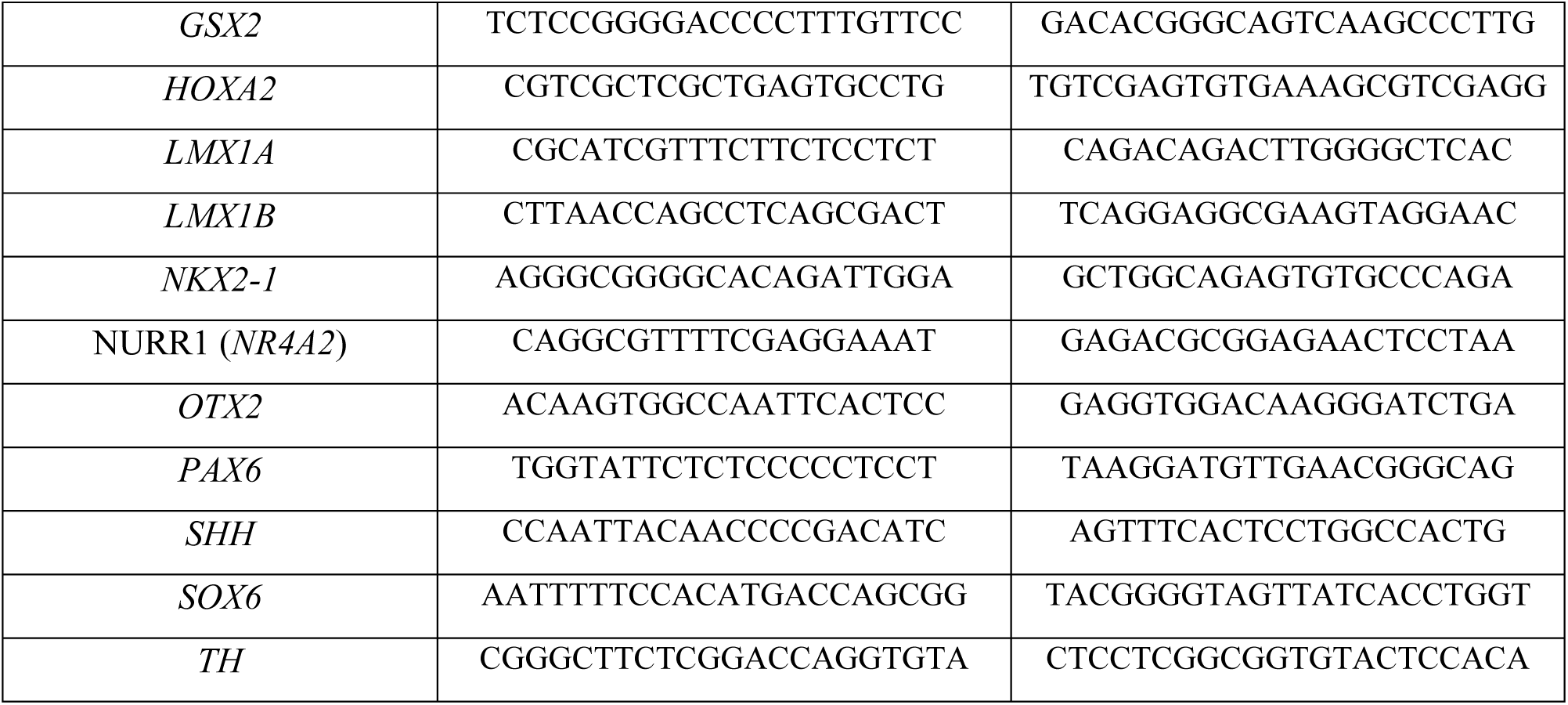
Sequence of qPCR primers.

### Organoid cryosectioning and immunofluorescence

Organoids were first gently dislodged from their respective connectoid compartments by aspirating them with a cut P1000 tip and then fixed in 4% paraformaldehyde (PFA) for 5 hours at room temperature or overnight at 4°C. After fixation, organoids were transferred into a 30% sucrose solution and left to sink overnight, at 4°C. The day after, organoids were transferred into a 1:1 mixture of 30% sucrose solution and OCT cryomount, overnight at 4°C. Finally, organoids were embedded in OCT cryomount within plastic molds, snap-frozen on dry ice, and sectioned at 20 µm thickness using a cryostat. For immunofluorescence staining, organoid sections were washed three times with 1× KPBS, then post-fixed in 4% PFA for 10 minutes at room temperature, followed by three additional washes. Slides were incubated in blocking solution (0.3% Triton X-100 and 5% donkey serum in 1× KPBS) for at least 1 hour. Primary antibodies (see Table 2) diluted in blocking solution were applied to slides incubated overnight at 4°C. After washing, sections were incubated with Alexa Fluor 488-, 594-, or 647-conjugated secondary antibodies (1:400; Jackson ImmunoResearch Laboratories) and DAPI (1:1000) in blocking solution for 1 hour at room temperature. Finally, sections were mounted on glass coverslips using FluorSave Reagent and imaged at least 24 hours later. For the 3D staining shown in Figs. 4A and EV4A, the same procedure was followed, except that blocking was performed overnight at room temperature, primary antibody incubation was extended to 48 hours, and secondary antibody incubation to 24 hours.

**Table 2.**
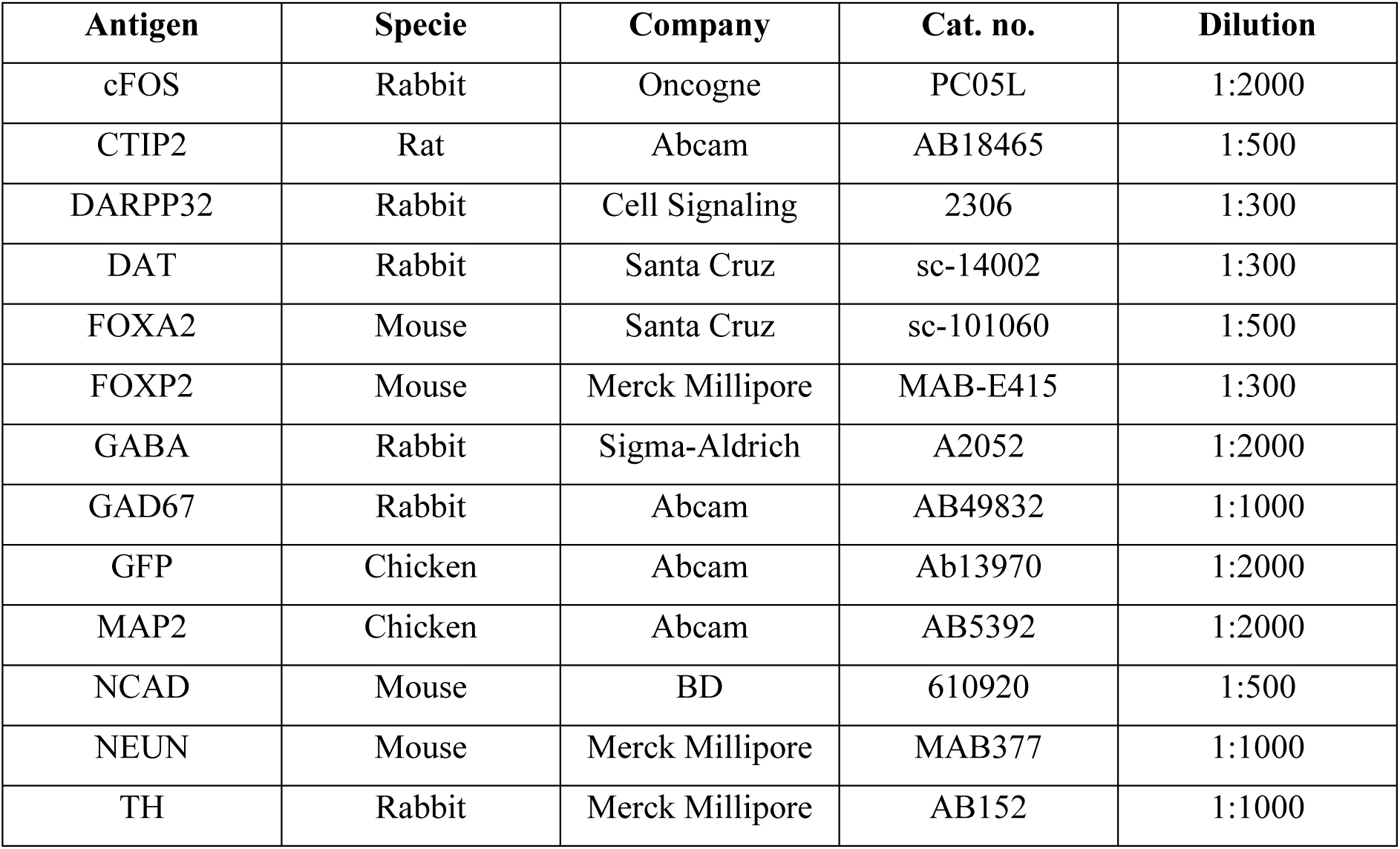
Primary antibodies.

### Single nuclei RNA sequencing sample preparation and raw data processing

Organoids were separated from the connectoid device using a P1000 pipette with cut tips, transferred to 1.5 mL Eppendorf tubes, and snap-frozen on dry ice. Nuclei were extracted and isolated from frozen pellets as described in Sozzi et al., STAR Protocols, 2025 (Sozzi *et al*, 2025). Samples were dissociated on ice in 1 mL nuclei lysis buffer (0.32 M sucrose, 5 mM CaCl_2_, 3 mM MgAc, 0.1 mM Na_2_EDTA, 10 mM Tris-HCl pH 8.0, 1 mM DTT, 0.1% Triton X) using glass Dounce homogenizers. Homogenates were centrifuged at 900 g for 15 min at 4 °C, and nuclei pellets resuspended in 300 μL wash buffer, composed of 0.1% BSA fraction V, 0.04 U/μL, and 0.02 U/μL RNase inhibitors (Ambion™, SUPERase In™, Invitrogen) in RNase-free PBS. Nuclei were purified by FANS on a FACSAria III (BD Biosciences, 100 μm nozzle) based on DRAQ7 (1:1000; D15106, ThermoFisher) staining and event size, with 12k nuclei per sample collected into BSA-precoated DNA LoBind tubes (Eppendorf) for snRNA-seq library preparation.

Single nuclei were loaded onto Chromium Single Cell 3′ Chips (10x Genomics, PN-120233, v3.1) with master mix and barcoded beads for encapsulation. Libraries were pooled, amplified, and then sequenced on an Illumina NovaSeq X in 2×100 bp paired-end mode. Demultiplexing was performed with bcl2fastq (v2.19), and raw transcriptome data was processed with cellranger (10x Genomics, v6.0) and aligned to the human genome assembly GRCh38 (2020-A). Genetic demultiplexing (RC17/H9 discrimination) was carried out with demuxlet (Kang *et al*, 2018), which assigns nuclei to their donor of origin based on inter-individual genetic variation. Reference genotype data were generated by whole-genome sequencing of individual hPSC lines using the Illumina DNA PCR-Free Prep Kit, sequenced on a NovaSeq X at ∼30× coverage. Data were processed with the Sarek pipeline (Garcia *et al*, 2020), and demuxlet (v1.0.5) was applied to BAM files from cellranger, including a third cell line (HS1001) as negative control.

### Bioinformatic analysis of single nuclei RNA sequencing data

Downstream processing was performed in R using the Seurat pipeline. During quality control, nuclei with a mitochondrial fraction greater than 1%, fewer than 500 detected features or with a H9 genetic background (STR-derived) were excluded from further analysis. Data integration across datasets was performed with the Harmony package (v1.2) to correct for batch effects. The resulting corrected coordinates were used for downstream projections (UMAP) and clustering, carried out using the Louvain algorithm, with the optimal resolution determined via the silhouette function from the cluster package (v2.1). Gene set enrichment analysis was conducted using the escape (v2.5) and clusterProfiler (v4.16) packages.

### Image acquisition and quantifications

Brightfield images were acquired using an Olympus CKX53 inverted microscope equipped with Olympus cellSens software (v2.3). Morphological assessments of developing vMB and STR organoids were performed every 2–3 days between day 5 and day 34 post-seeding (n = 5 organoids per condition per time point) and included measurements of maximum cross-sectional area and circularity, defined as 4*π* × area / perimeter^2^. Fluorescent staining images were captured using a Leica DMI6000B widefield microscope equipped with Leica LAS X acquisition software. Time-lapse imaging of dopaminergic neuron outgrowth and degeneration was performed on a Leica TCS SP8 confocal microscope over a period of 10 days, using the connectoid device as a reference for image alignment. Images were acquired every 12 hours during axonal growth and every 48 hours during 6-OHDA treatment using consistent acquisition parameters (1024 × 1024 px tile scan, 10x air objective, 150 µm z-stack, 21 z-planes, pinhole 2 AU). Quantifications were performed in ImageJ2 (v2.16) after applying a Gaussian blur filter (σ = 1) and automatic thresholding using the Triangle method on manually defined regions of interest (ROIs) corresponding to either the vMB compartment or the connectoid channel. Neurite length over time was measured by tracing neurites on maximum-intensity projection images using the Single Neurite Tracer (SNT) tool within the Neuroanatomy plugin in ImageJ2. All fluorescence and brightfield images were processed using Adobe Photoshop CC 2020, with identical adjustments applied uniformly across the entire image.

### Rabies-based retrograde tracing

The TVA receptor and the corresponding GP-depleted control construct were obtained from Addgene (plasmid IDs: 30195 and 30456, respectively) and integrated in the developing STR organoids at day 7 (MOI 2). EnvA-pseudotyped, GP-deprived rabies virus (EnvA-ΔG-RV) was produced as previously described in Grealish et al. 2015 (Grealish *et al*, 2015) and applied to cultures at a 0.5% v/v dilution in terminal differentiation medium at day 63 of differentiation for 48 hours. Confocal imaging was performed using a Leica TCS SP8 microscope equipped with LAS X acquisition software.

### Pharmacological treatments and optogenetic stimulation

To induce degeneration of DAergic fibers, vMB-STR connectoids were treated with 400 μM 6-hydroxydopamine (6-OHDA) for 48 hours at day 60 of differentiation. Pharmacological activation of DA synapses was achieved by exposing connectoids to 20 μM D-amphetamine (Apoteksbolaget, Sweden) for 2 hours prior to fixation. For optogenetic experiments, TH-Cre–derived vMB organoids were transduced at day 7 (MOI 2) with lentiviral particles carrying a YFP::flexChR2 construct. ChR2 stimulation was performed using 570 nm blue light delivered through an optic fiber on the vMB compartment for 10 minutes, and STR organoids were fixed 1.5 hours later. cFOS quantifications were performed using ImageJ2 by thresholding DAPI-stained nuclei with the automatic Otsu algorithm, followed by particle counting between 0.3x and 3x the median particle area, estimated from 20 nuclei per image (n = 3 organoids per group; average nuclei count per organoid = 1089 ± 85). cFOS-positive nuclei were defined as those with an average fluorescence intensity at least two standard deviations above the mean background signal.

### Dopamine detection

For dopamine release assessment, the media was removed, and the connectoids were washed 3 times with 500 μL of HBSS. They were then incubated in 300-500 μL of HBSS at 37°C for one hour. Following the incubation, the supernatant was harvested into Eppendorf tubes. Samples were snap-frozen immediately after collection and stored at −80 °C until analysis. HEK-293 Flp-In T-REx cells engineered to stably express the fluorescent GPCR-based dopamine sensor GRAB_DA2M_ (Klein Herenbrink *et al*., 2022; Sun *et al*., 2018) were maintained in DMEM containing 10% FBS, 15 µg/ml blasticidin, and 200 µg/ml hygromycin B. One day prior to imaging, the cells were plated into poly-L-ornithine–coated imaging chambers (18-well, ibidi) and sensor expression was induced with 1 µg/ml tetracycline. Live imaging of GRAB_DA2M_ signals was carried out on a Leica widefield microscope using a 20x objective. Three images were taken per well: baseline fluorescence, image after the addition of sample collected from brain organoids, and a maximum fluorescence response after the addition of 10 μM DA solution. Mean fluorescence intensity was extracted for each image using ImageJ (NIH) and the response was reported as %max.

## Acknowledgements

We would like to thank Alessandro Fiorenzano and Petter Storm for their scientific inputs and support, and Ulla Jarl, Jenny G Johansson, and Bengt Mattsson for excellent technical assistance. We gratefully acknowledge funding from the HORIZON-EIC-2021-PATHFINDEROPEN-01 project 101047177 OpenMIND, HORIZON-ERC-2024-SYG, project 101167102 Custom-Made, Swedish Research Council (M.P. VR rådsprofessur n°2021-00661, 3R-VR n°2021-02967), Swedish Parkinson Foundation (Parkinsonfonden, 1675/25), Swedish Brain Foundation, (Hjärnfonden, FO2024-01), Mats Paulsson Foundation, Konung Gustaf V:s och Drottning Victorias Frimurarestiftelsen, the Strategic Research Areas at Lund University MultiPark (Multidisciplinary research in Parkinson’s disease) and StemTherapy/Lund Stem Cell Center. SC was supported by the Royal Physiographic Society in Lund (45540). The authors would also like to acknowledge Clinical Genomics Lund, SciLifeLab and Center for Translational Genomics (CTG), Lund University, for providing sequencing service.

## Author contributions

JE and MP conceived the study, supervised the project and interpreted data. ES, SC, AB, GS, SM, ST, AH, GR, JK designed and/or performed the experiments and interpreted data. ES, SC, and AB prepared figures. ES, SC, AB and MP wrote the manuscript with input from all authors. All authors read and approved the final version.

## Data availability

Raw single-nucleus RNA sequencing data generated in this study are publicly available at the Gene Expression Omnibus (GEO) under accession number GSE312520. The analysis code related to this work is available from the leading author upon request.

## Expanded View Figure legends

**Figure EV1:**
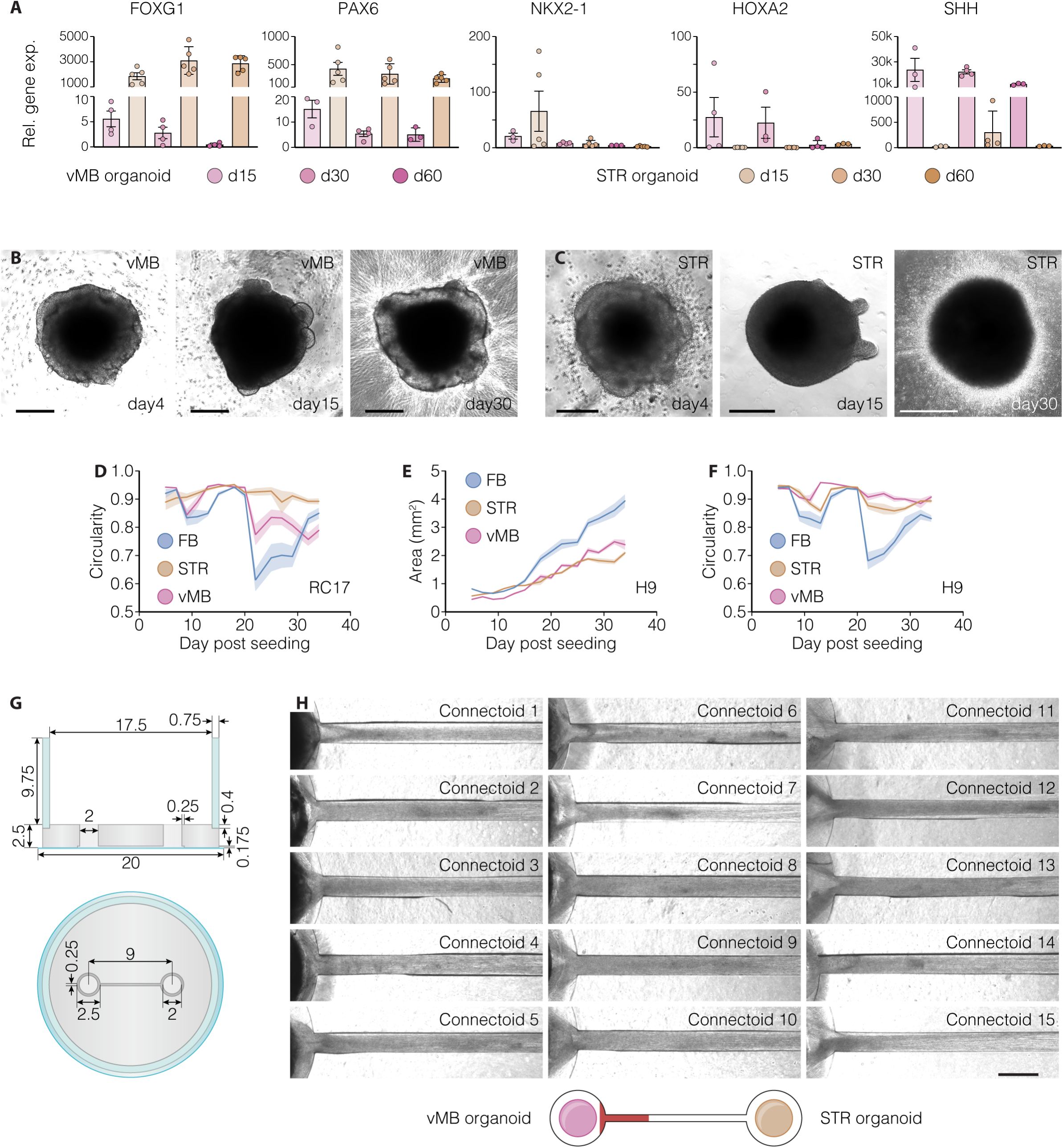
A) Relative expression levels of key regional markers in vMB and STR organoids at days 15, 30, and 60 of differentiation. B, C) Representative brightfield images of developing vMB (B) and STR (C) organoids at days 4, 15, and 30 of differentiation. Scale bars: 200 μm (day 4), 300 μm (day 15), 500 μm (day 30). D–F) Morphological analysis of vMB, STR, and unguided FB organoids over time (days 5–34 post-seeding), showing circularity in RC17-derived organoids (D) and maximum cross-sectional area (E) and circularity (F) in H9-derived organoids. G) Schematic side and top views of the connectoid device, with all dimensions indicated (in mm). H) Brightfield images of 15 individual connectoids generated within the same batch at day 40, illustrating the reproducible formation of axonal bundles within the connectoid channel. The red-highlighted region in the schematic below indicates the area imaged. Scale bar, 500 μm.

**Figure EV2:**
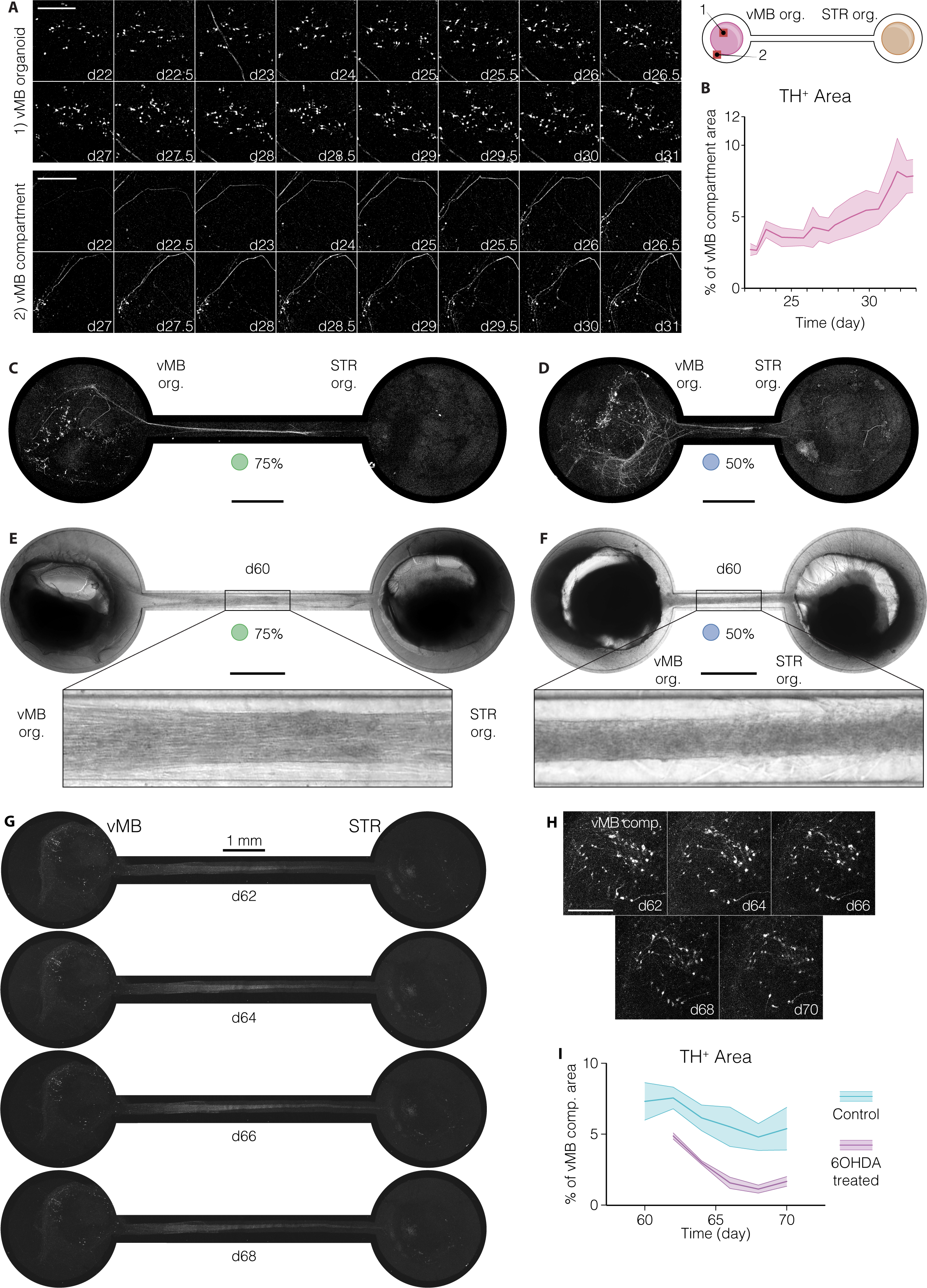
A) Time-lapse imaging of GFP⁺ cells in vMB organoids (1) and the vMB compartment (2) of vMB–STR connectoids between days 22 and 32. Red boxes in the connectoid schematic indicate the corresponding fields of view. Scale bars, 200 μm. B) Quantification of GFP/TH⁺ area in the vMB compartment over time, presented as mean ± SEM. C, D) Maximum intensity projection confocal images of vMB–STR connectoids with 75% (C) and 50% (D) channel length at day 32. Scale bars, 1 mm. E, F) Representative brightfield images of day 60 vMB–STR connectoids with 75% (E) and 50% (F) channel length compared to the original 9 mm design, highlighting the formation of the axonal bundle. Scale bars, 1 mm. G, H) Overview (G) and high-magnification (H) images of control vMB–STR connectoids between days 62 and 70, showing GFP/TH⁺ long-range projecting DA neurons over time. I) Quantification of TH⁺ area in the vMB compartment of control and 6-OHDA-treated vMB–STR connectoids between days 60 and 70 (Friedman test, p = 0.0006).

**Figure EV 3:**
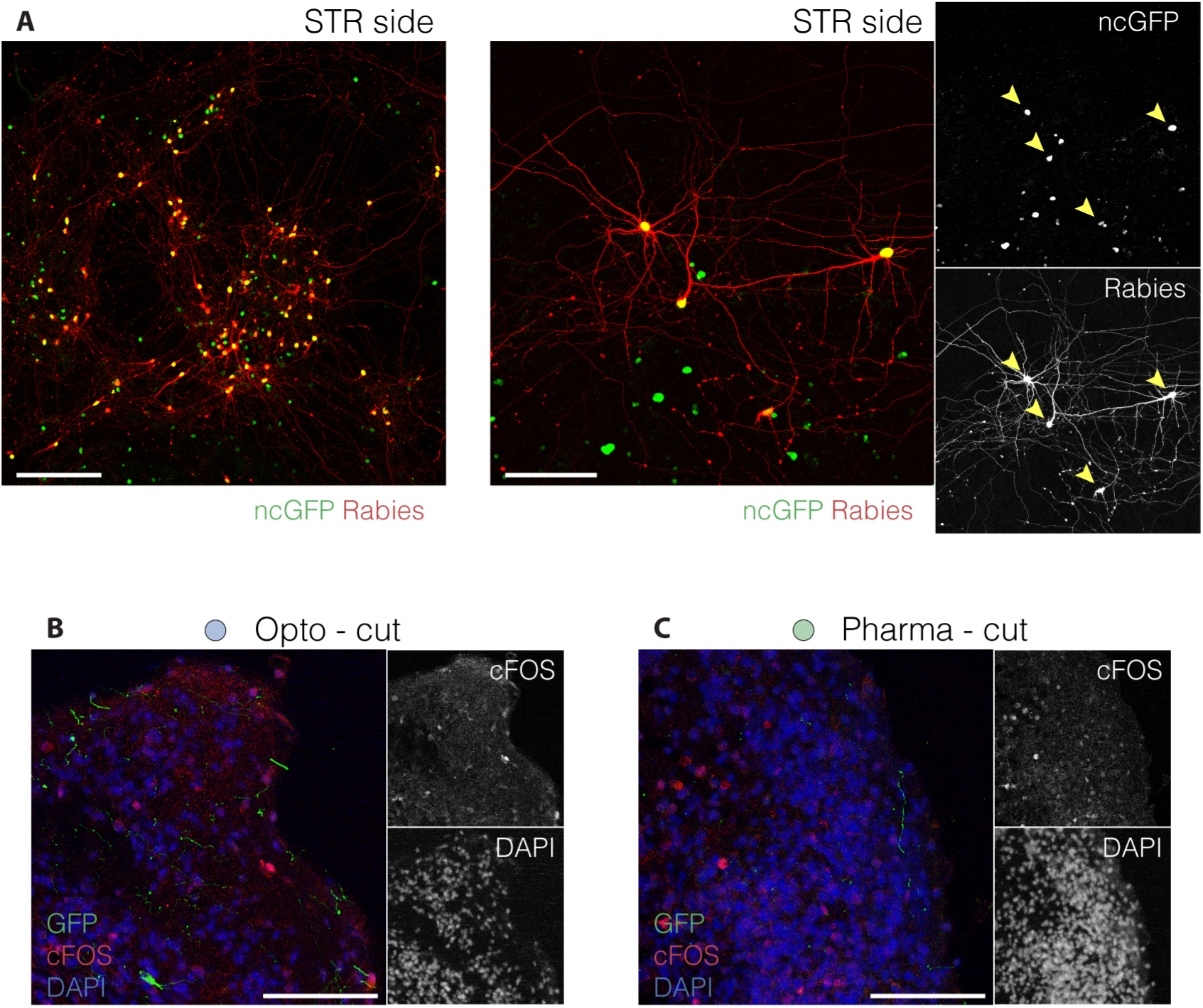
A) Low- (left) and high-magnification (right) confocal images of control STR organoids within vMB–STR connectoids transduced with a GP-negative helper tracing vector. The yellow arrowhead indicates mCherry⁺ cell bodies, all co-expressing nuclear GFP from the tracing vector. Scale bars: 200 μm (left); 100 μm (right). B,C) Confocal images of GFP, cFOS, and DAPI immunostaining in the STR compartment of severed connectoids following stimulation with 470 nm blue light (B) or 20 μM D-amphetamine (C) at day 95. Corresponding single-channel images of cFOS and DAPI used for quantification are shown alongside. Scale bars, 100 μm.

**Figure EV 4:**
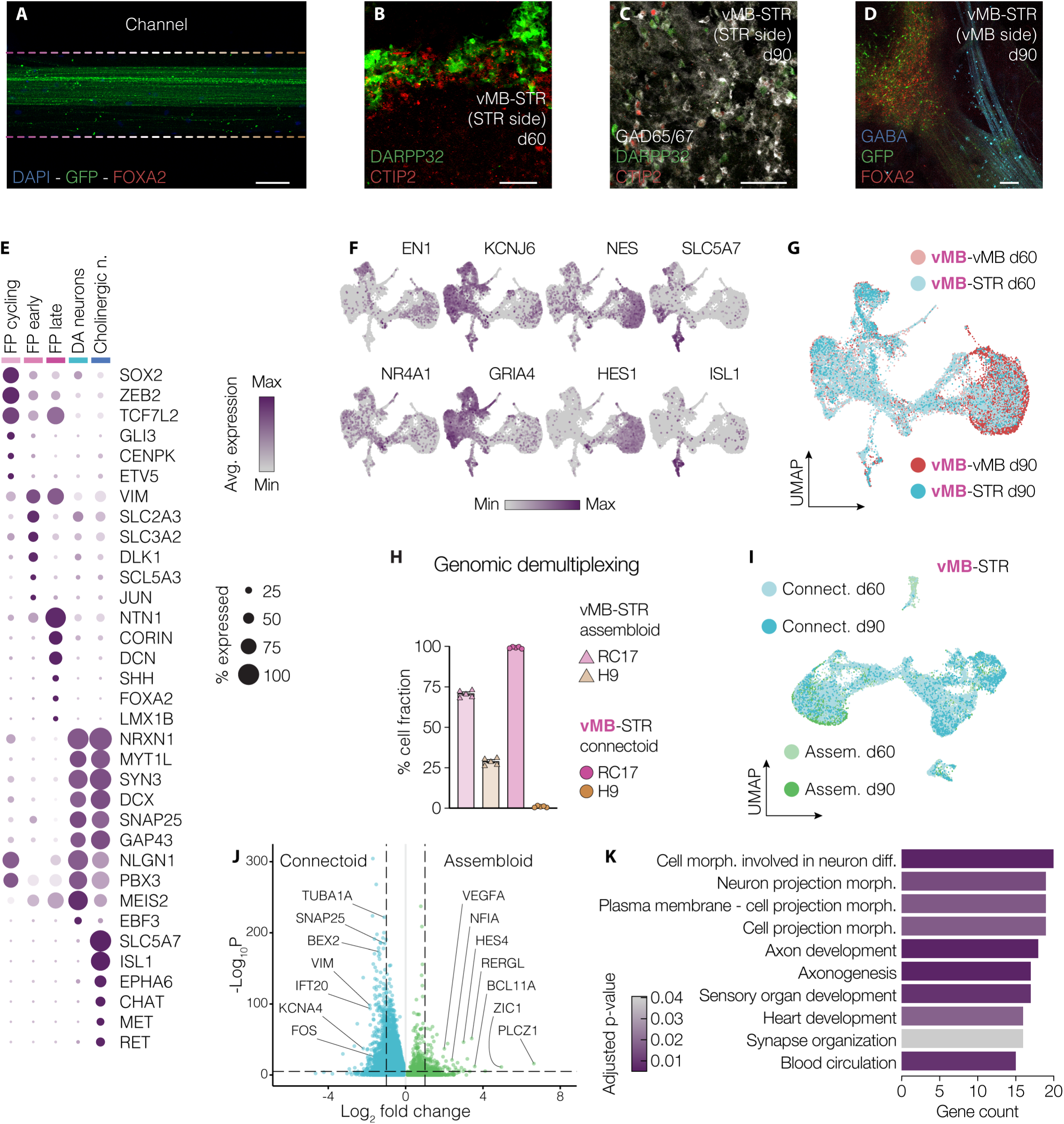
A) Immunohistochemistry for GFP and FOXA2 in the intact channel of vMB–STR connectoids at day 30. Nuclei are stained with DAPI. Scale bar, 100 μm. B, C) Fluorescence images of vMB–STR connectoids at day 60 (B) and day 90 (C), showing expression of DARPP32 and CTIP2 (B), and GAD65/67, DARPP32, and CTIP2 (C) in the STR compartment. Scale bars, 100 μm. D) Immunofluorescence staining for GFP, GABA, and FOXA2 in the vMB compartment of vMB–STR connectoids at day 90. Scale bar, 100 μm. E) Dot plot showing expression levels of selected marker genes across clusters in the vMB compartment of connectoids between days 60 and 90. N., neurons. F) Feature plots showing the distribution of cells expressing selected markers in the vMB compartment dataset. G) UMAP representation of the vMB compartment dataset derived from vMB–vMB and vMB–STR connectoids at days 60 and 90. H) Bar plot showing the relative cell fractions generated from RC17 and H9 hPSC lines in vMB–STR assembloids (triangles) and in the vMB compartment of vMB–STR connectoids (circles; n = 5). I) UMAP plot of the vMB component of vMB–STR assembloids and connectoids at days 60 and 90. J) Volcano plot showing differentially expressed genes between DA neurons derived from vMB–STR connectoids and assembloids. K) Enrichment analysis of the top ten gene ontology biological process terms associated with upregulated genes in connectoid-derived DA neurons compared to assembloid-derived DA neurons. Morph., morphogenesis.

